# Violet light modulates the central nervous system to regulate memory and mood

**DOI:** 10.1101/2021.11.02.466604

**Authors:** Nobunari Sasaki, Pooja Gusain, Motoshi Hayano, Tetsuro Sugaya, Naoya Tonegawa, Yusuke Hatanaka, Risako Tamura, Kei Okuyama, Hideto Osada, Norimitsu Ban, Yasue Mitsukura, Richard A. Lang, Masaru Mimura, Kazuo Tsubota

## Abstract

Light stimuli from the external environment serves as a signal. Photoreceptors receive photons at the outer nuclear layer of the retina. Non-visual photoreceptors, such as opsin5 (also known as OPN5 or neuropsin), are expressed in the retinal ganglion cells (RGCs) and hypothalamus to regulate the circadian cycle and body temperature. Here, we show that violet light (VL) stimuli received by OPN5-positive RGCs are transmitted to the habenula brain region. VL improves memory in aged mice and simultaneously increases neural architecture-related genes such as oligodendrocyte-related genes in the hippocampus. In addition, VL improves depressive-like behaviors in the social defeat stress model in an OPN5 dependent manner. Following VL exposure, cFos activation is observed at the nucleus accumbens (NAc) and the paraventricular thalamic nucleus (PVT). Taken together, the results indicate that violet light modulates brain function such as memory and mood by transmitting the signal from RGCs to the habenula region in the brain.

## Main

Sunlight has been utilized evolutionarily as an energy source to produce ATP or as a signal for biological control since before organisms developed eyes^1^. In vertebrates, visible light from 360 nm to 700 nm serves not only a visual function but also a non-visual function such as blue light stimulation regulates sleep, memory, and emotion by melanopsin (OPN4) in intrinsically photosensitive retinal ganglion cells (ipRGCs)^2,3^. OPN4 is a non-visual opsin and sensitive to blue light (λmax = 480nm)^4^. OPN4-positive ipRGCs are further classified into *Brn3b*(+) and *Brn3b*(-) ipRGCs^5^. *Brn3b*(+) ipRGCs directly project the light signal to the perihabenular nucleus (PHb) or indirectly project to the lateral habenula (LHb) via the ventral lateral geniculate nucleus-intergeniculate leaflet (vLGN–IGL) to regulate mood and depression^6,7^. On the other hand, *Brn3b*(-) ipRGCs function to establish a IGL^NPY^-SCN (suprachiasmatic nucleus) circuit controlling nonphotic entrainment to food^6^. The signal from *Brn3b*(+) ipRGCs to the SCN is also known to regulate sleep and memory function^2,3^.

Neuropsin (OPN5) is the most recently identified non-visual opsin^8^. OPN5 is a Gi-type G protein-coupled receptor (GPCR) proved to be a visible violet light (VL) responsive opsin (λmax = 380nm)^9^. Mammalian OPN5 contributes to photoentrainment to light/dark (LD) cycle and is also implicated in the local circadian photoentrainment in retina, cornea and skin^10–12^. Additionally, OPN5 expression in warm-sensing neurons in the hypothalamic preoptic area (POA) was shown to regulate VL-evoked suppression of thermogenesis in brown adipose tissue (BAT)^13^.

Although OPN5 is expressed in RGCs and regulates the timing of hyaloid vascular regression in response to VL^14^, its expression does not overlap with OPN4 indicating that OPN5-expressing RGCs have distinct functions from OPN4-expressing RGCs. In addition, it remains unknown where OPN5-expressing RGCs project to and whether the VL-OPN5 axis is involved in the regulation of brain functions such as learning and mood.

In the current study, tracer experiments identify OPN5-expressing RGCs projecting to the habenula region. We show that cognitive function is increased by VL exposure in aged mice in which myelinating oligodendrocyte related genes are increased and neural excitation-related genes are decreased in the hippocampus while simultaneously increasing cFos-positive cells. In addition, VL improves depressive behavior induced by social defeat stress (SDS) and cFos-positive cells are increased at nucleus accumbens (NAc) and paraventricular thalamic nucleus (PVT). We demonstrate novel functions of VL and potential therapeutic approaches using VL for dementia and depression.

## Results

### Identification of subtypes of *Opn5*-expressing RGCs

Single RNA sequencing (RNA-seq) revealed that RGCs can be classified into 46 subtypes based on gene expression^15^. Furthermore, RGC subtypes can be classified according to their distinct electrophysiological responses to light stimuli. To determine the subtype of *Opn5*-expressing RGCs (OPN5-RGCs), we used a *Opn5^cre^; Ai14* combination mouse, in which tdTomato is expressed in the *Opn5* lineage^14^. As previously reported, tdTomato was observed in the RGC layer of the retina (Fig. 1a). Immunolabeling of *Opn5^cre^; Ai14* mice showed that 99.4% of tdTomato-labeled cells were also positive for *RBPMS*, an RNA-binding protein and a pan-RGC marker. This indicates that *Opn5*-expressing cells are RGCs (Fig. 1b). Consistent with previous reports^14^, *Opn5*-expressing RGCs were not co-labeled with antibodies to melanopsin/OPN4. In addition, 94.3% of tdTomato-labeled cells co-label with antibodies to Brn3a, a POU domain transcription factor family member and a negative marker of ipRGCs. This further indicates that OPN5-RGCs are distinct from ipRGCs (Fig. 1c). Approximately 26% of ipRGCs release γ-aminobutyric acid (GABA, an inhibitory neurotransmitter) to regulate the pupillary light reflex (PLR) and circadian behavior^16^. The GABA subpopulation of ipRGCs is marked by GAD65, an enzyme involved in the biosynthesis of GABA. *Opn5* lineage tdTomato-labeled cells were not co-labeled with GAD65 (Fig. 1d). Axons from ipRGCs project to the SCN, LHb and PHb. To assess projections from OPN5-RGCs, we examined tdTomato-labeled nerve fibers in *Opn5^cre^; Ai14* mice. tdTomato-labeled nerve fibers were observed in the optic tract and dorsal lateral geniculate nucleus (dLGN) as previously reported, but not in the SCN (Fig.1e, f). Taken together, these results suggest that OPN5-positive RGCs, unlike melanopsin-positive ipRGCs, are conventional RGCs that project to the brain via the optic nerve.

**Fig. 1.**
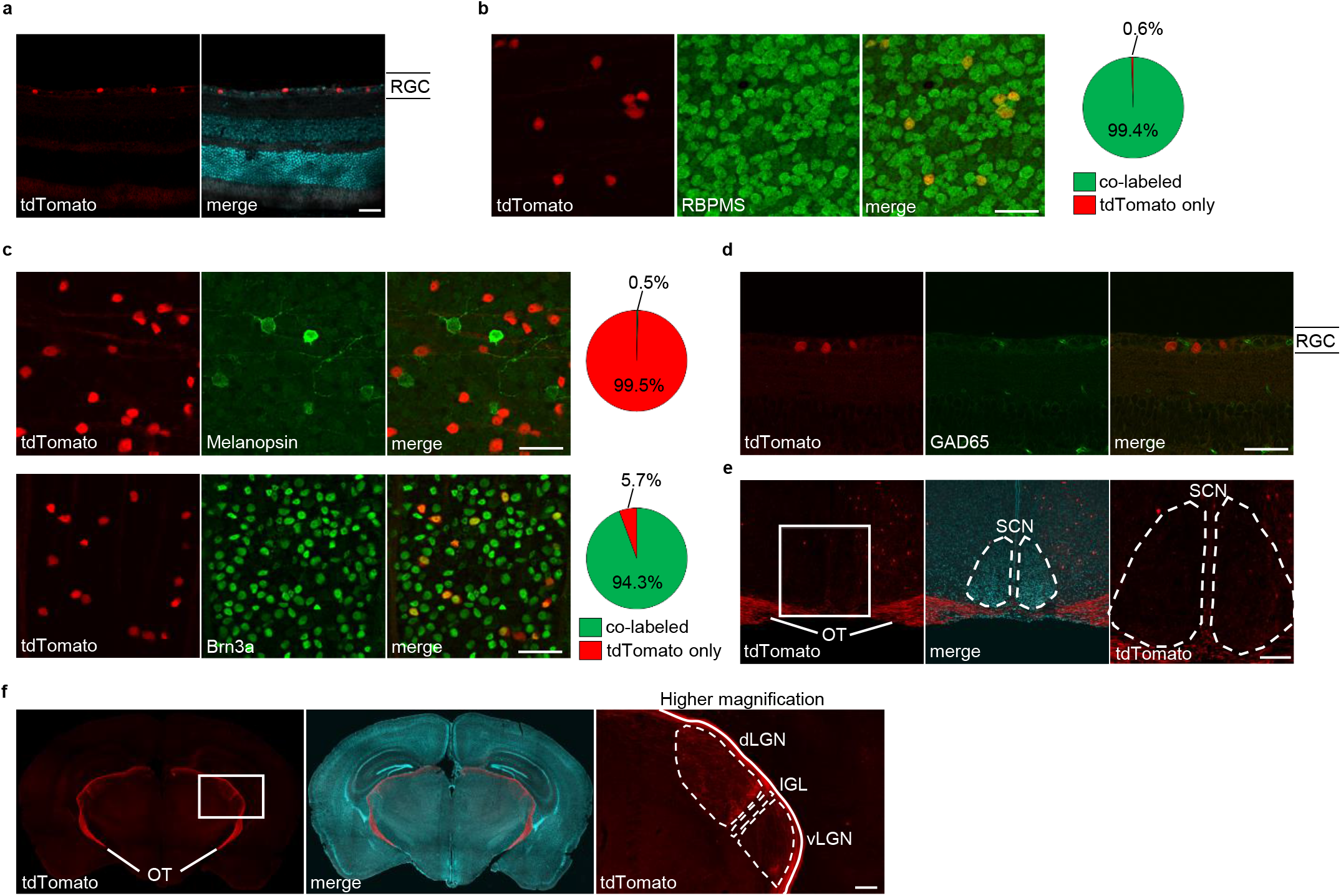
Identification of *Opn5*-expressing RGCs and nerve fibers in brain. **a**, Frozen section images of *Opn5^cre^; Ai14* mouse retina; merged images show the overlaid image of tdTomato and DAPI staining. RGC, retinal ganglion cell layer. **b**, RBPMS immunostaining image of retinal whole mount of *Opn5^cre^; Ai14* mouse. Pie charts show the percentage of cells that are single stained for tdTomato and those that are co-stained for tdTomato and RBPMS. (n=3) **c**, Melanopsin, Brn3a immunostaining image of *Opn5^cre^; Ai14* mouse retinal whole mount. The pie chart shows the percentage of cells that are single stained for tdTomato and those that are co-stained for tdTomato and Melanopsin or Brn3a (n=3). **d**, GAD65 immunostaining images of *Opn5^cre^; Ai14* mouse retinal cryosections. **e**, *Opn5^cre^; Ai14* mouse brain frozen section images. Merge image shows the overlaid image of tdTomato and DAPI staining. The right panel shows a magnified image of the area enclosed by the white square in the left panel. OT, optic tract; SCN, suprachiasmatic nucleus. **f**, Frozen section image of *Opn5^cre^; Ai14* mouse brain. Merge image shows the overlaid image of tdTomato and DAPI staining. The right panel shows a magnified image of the area enclosed by the white square in the left panel. dLGN, dorsal lateral geniculate nucleus; vLGN, ventral lateral geniculate nucleus; IGL, intergeniculate leaflet, Scale bar (**a**,**b**,**c**, and **d**) 40 μm, (**e**) 100 μm and (**f**) 200 μm.

### Tracking of nerve fibers from *Opn5*-expressing RGCs

To assess the brain regions to which OPN5-RGCs project, we performed tracing experiments using an adeno-associated virus, AAV-DIO (double-floxed inverted open reading frame)-WGA, that expresses wheat germ agglutinin (WGA) in a Cre-dependent manner (Fig. 2a). The WGA protein is expected to be transported across synapses in an anterograde manner^17,18^. Whole-mount observation of the retinas of *Opn5^cre^; Ai14* mice injected intravitreally with AAV-DIO-WGA showed that WGA was expressed in tdTomato-positive RGCs (Fig. 2b). Furthermore, immunostaining of retinas from *Opn5^cre^* mice injected intravitreally with AAV-DIO-WGA showed that WGA was expressed only in the RGC layer. These data indicate that WGA expression is restricted to OPN5-RGCs (Fig. 2c). We then examined the localization of WGA in the brain of *Opn5^cre^* mice after injecting AAV-DIO-WGA. WGA expression was detected in the habenula region, especially in the medial habenula (MHb) (Fig. 2d). Taken together, these data indicate that the input from OPN5-RGCs projects to the habenula.

**Fig. 2.**
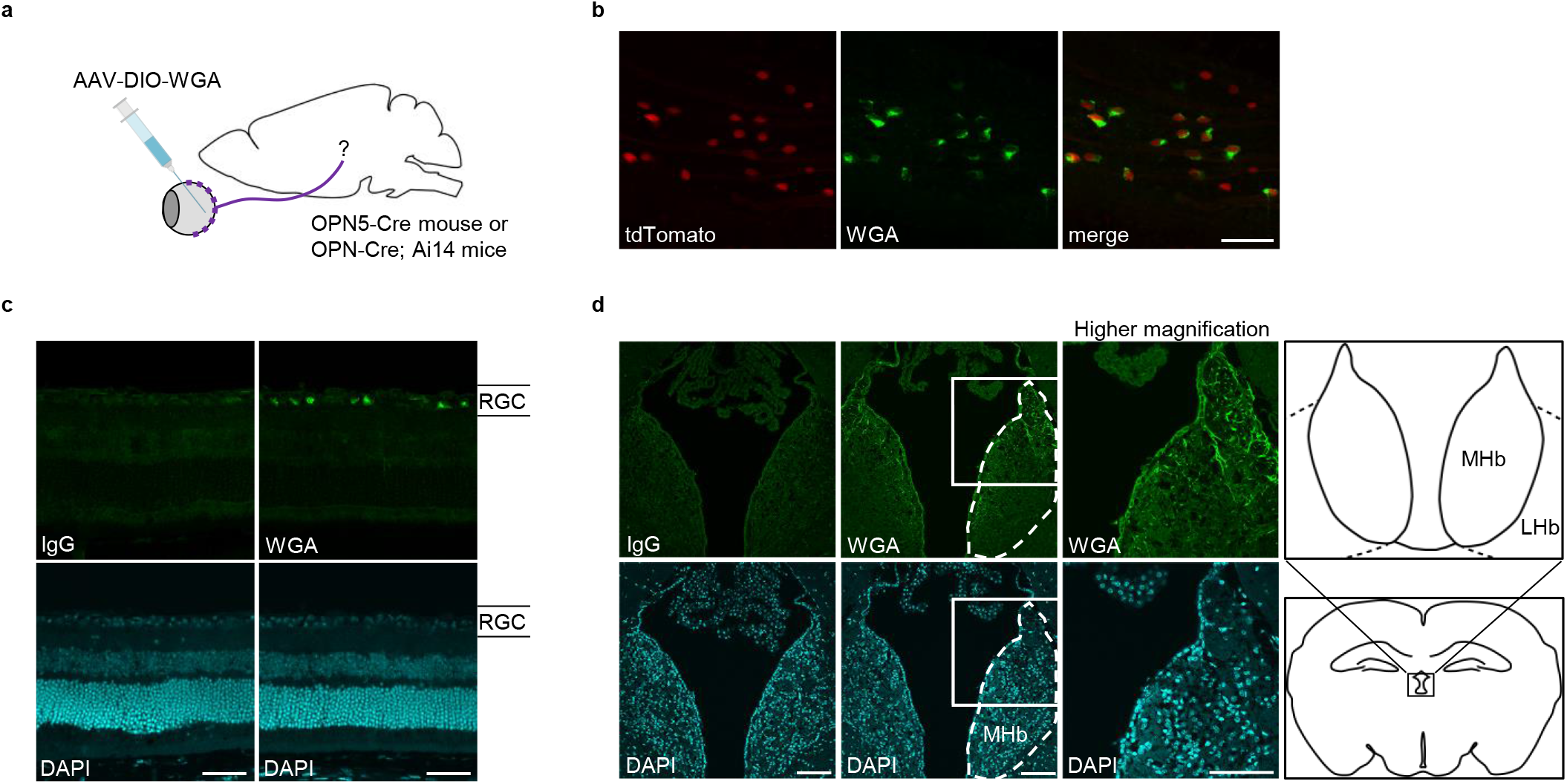
Tracking of retinal fibers from *Opn5*-expressing RGCs. **a**, Schematic of neuronal tracing with AAV-DIO-WGA, which was injected into the vitreous of *Opn5^cre^* or *Opn5^cre^; Ai14* mice to selectively express WGA protein in OPN5-RGCs. **b**, Whole-mount immunostaining images of retinas from *Opn5^cre^; Ai14* mice injected with AAV-DIO-WGA. **c**, Immunostained images of frozen sections of *Opn5^cre^* mouse retina injected with AAV-DIO-WGA. WGA (green, upper panel) and nuclei (blue, lower panel) are labeled. **d**, Expression of WGA in the habenula of mouse brain. The center-right panel shows an enlarged image of the region indicated by the white square in the center-left panel. The right panel shows a schematic of the habenula and coronal sections of the brain. Representative images are shown in (**c**) and (**d**) (n=3). Scale bar is (**b**) 40 μm, (**c**) 50 μm, and (**d**) 100 μm.

### Regulation of learning and memory by VL exposure

To examine the effects of VL stimulation on cognitive function, aged mice were exposed to VL for 2 h per day (ZT2-ZT4) (Fig. 3a and Extended Data Fig. 1a, b). The *Opn5* gene expression in the aged mice retina was comparable to that in young mice (Extended Data Fig. 2a). After 7 weeks of VL exposure, cognitive function was assessed using the contextual fear conditioning (CFC) test, a behavioral experiment that evaluates contextual fear memory for electrical foot shocks (Fig. 3a). Although the percentage of freezing immediately after electroshock did not differ between the mice exposed to white light and VL, the contextual freezing behavior one day later was significantly increased in the mice exposed to VL compared to white light, indicating that the VL stimulation has a positive effect on cognitive function (Fig. 3b). To further assess cognitive function as working memory, we performed the Barnes maze test, a behavioral test commonly used to evaluate spatial learning and memory. Mice were subjected to Barns maze test for a total duration of 8 days including 6 days of adaptive training, followed by a probe trial to assess memory retention. The time spent around the target hole was significantly increased in the group exposed to VL (Fig. 3c). Anxiety-related behaviors and locomotion were assessed using the open field test (OFT). VL exposure had no effect on the duration of time spent in the central region or total movement distance in the field (Fig. 3d, e). Then, we determined whether neuronal activation is induced by VL in the brain. Immunostaining of cFos in frozen sections of the brains of aged mice exposed to VL revealed a significant increase in the number of cells expressing cFos in the hippocampal CA1 region, dentate gyrus, and LHb (Fig. 3f, g and Extended Data Fig. 3a-d). These results suggest that VL stimulation impacts the age-dependent decline of cognitive function.

**Fig. 3.**
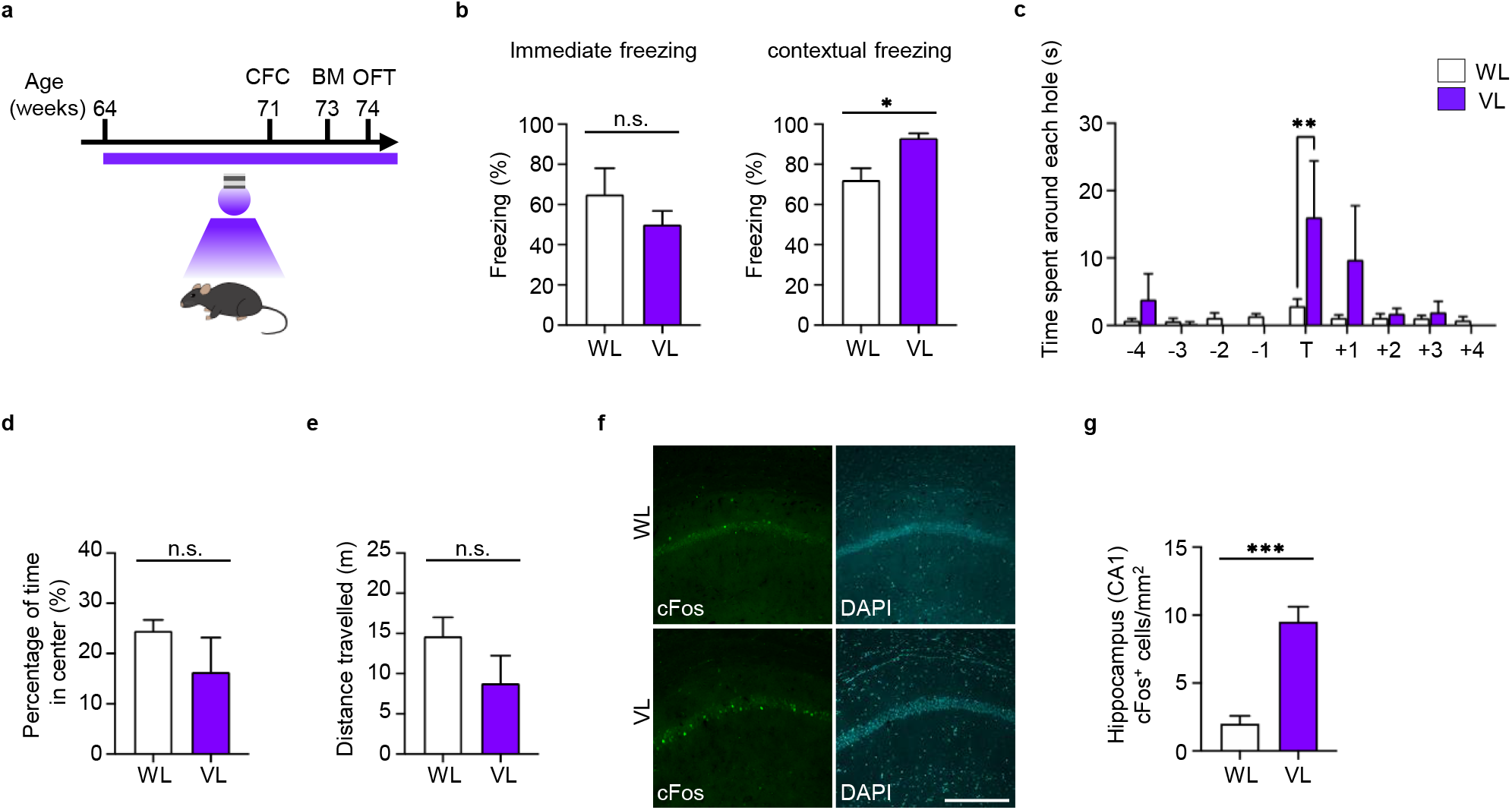
VL exposure improves memory function. **a**, Schematic of the experimental schedule. During the period shown in the figure, mice were exposed to VL for 2 h per day from 10 am to 12 pm. **b**, Freezing rate immediately after electroshock (left; immediate freezing) and 1 day later (right; contextual freezing) in the fear conditioning test. *:p=0.0198, n.s., not significant, Student’s t-test; VL (n=4), WL (n=5). **c**, Time spent around each hole in the probe test 8 days after the final training of the Barnes maze test. Holes were located clockwise from the target hole. **:p=0.0045, Two-way ANOVA analysis followed by Bonferroni test; VL (n=3), WL (n=4). Percentage of time spent in the inner area (**d**) and total distance traveled (**e**) in the open field test. Student’s t-test; VL (n=4), WL (n=5). **f**, Expression of cFos in the mouse brain exposed to VL. Frozen sections of mouse brains irradiated with white light (upper panel) or VL (lower panel) were immunostained with cFos and DAPI. **g,**Quantitative analysis of cFos positive cells in the hippocampus. ***p=0.0001, Student’s t-test; WL (n=3) and VL (n=3). Scale bar 50 μm. Bar graphs show mean±s.e.m.

### VL exposure changes the gene expression in hippocampus

To assess the effect of VL stimulation on the gene expression profile in the brain, we exposed aged mice to VL followed by performing RNA-seq in hippocampus. We identified 404 differentially expressed genes (DEGs), including 231 up-regulated and 173 down-regulated genes (Fig. 4a and Extended Data Fig. 4a). Gene Ontology (GO) analysis by classifying DEGs into up-regulated and down-regulated genes showed that VL stimulation increased expression of genes involved in neural circuit formation, such as ensheathment of neurons, axon ensheathment, and myelination (Fig. 4b). Glia such as astrocytes, microglia, and oligodendrocytes have a crucial role for synaptic maintenance and the enhancement of neurotransmission^19^. It is notable that glial cell development, especially oligodendrocyte differentiation, is included in the clusters of up-regulated genes. In contrast, VL stimulation suppressed expression of genes involved in neural excitation and membranes, including genes involved in the synapse, neuron projection, synaptic membrane, and intrinsic component of plasma membrane (Fig. 4c). VL-exposed aged mice show up-regulation of genes associated with oligodendrocyte differentiation. We hypothesized that VL improved cognitive function in aged mice by up-regulation of genes involved in the differentiation of oligodendrocytes. Specifically, genes up-regulated by VL play a crucial role in differentiation of pre-oligodendrocytes (pre-OL) to mature oligodendrocytes. This includes genes involved in myelin maturation such as *Sox10, Cnp, Mag, Mbp*, and *Mobp* (Fig. 4d, e). Furthermore, VL increased the expression of *Myrf*, a known Sox10 co-factor that induces oligodendrocyte differentiation genes. VL also induces other oligodendrocyte related genes including *Pou3f1* and *Transferrin* (*Trf*) (Fig. 4d). Neurotransmission and synapse-related genes such as *Atp2b4, Cadm1, Camk2d, Dnm3*, *Gap43* and *Gria4* were suppressed by VL in aged mice (Fig. 4f). Together, these data indicate that VL stimulation modifies the gene expression involved in glia differentiation and neural excitation.

**Fig. 4.**
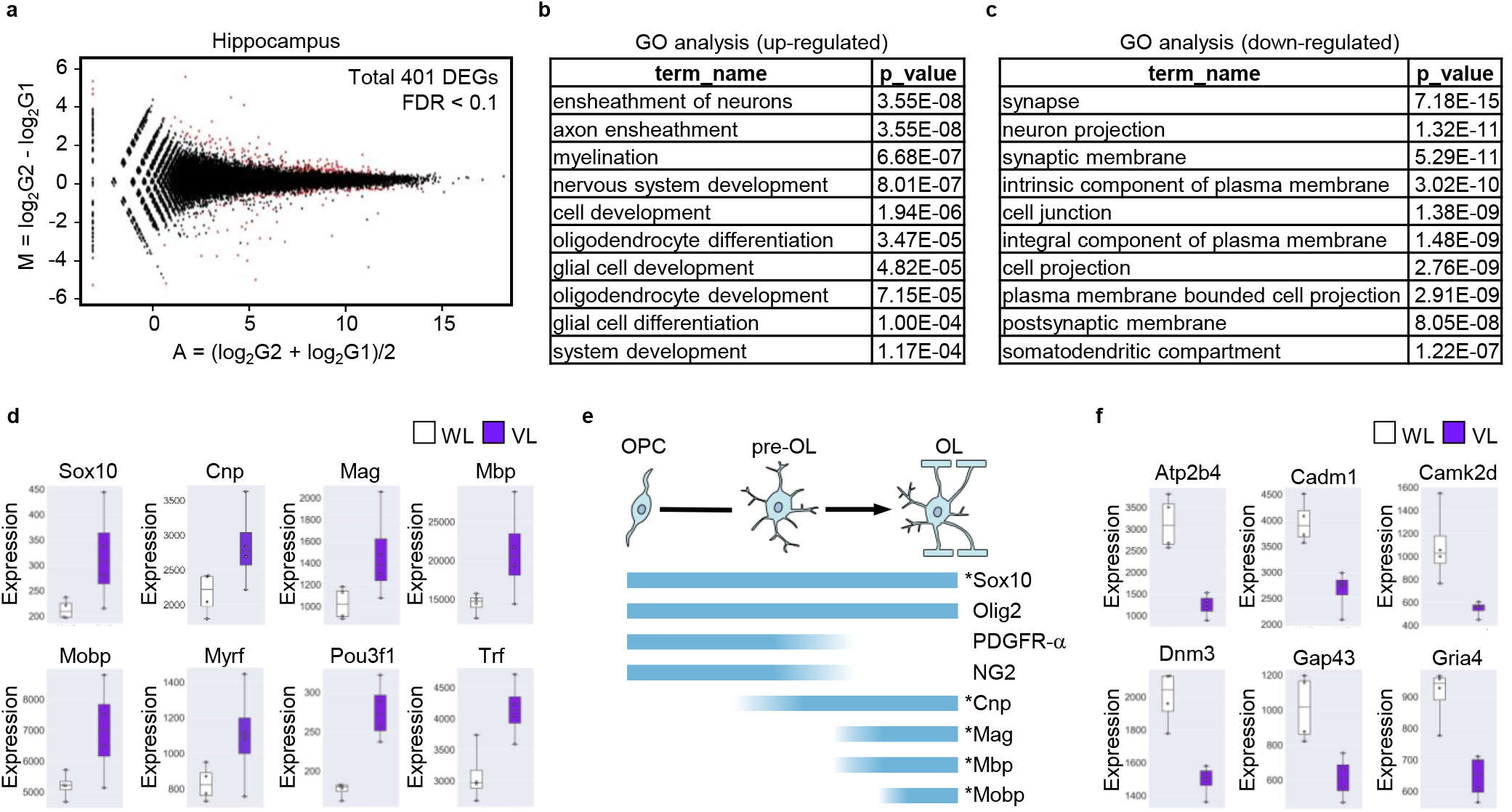
VL modulates gene expression in hippocampus. **a**, MA plot of changes in gene expression in the hippocampus of aged mice exposed to white light and VL; differentially expressed genes (DEGs) are shown in red (FDR<0.1). **b**, Gene ontology analysis using the genes whose expression was upregulated in the hippocampus of mice exposed to VL (FDR<0.1). **c**, Gene ontology analysis using genes whose expression was downregulated in the hippocampus of mice exposed to VL (FDR<0.1). **d**, Genes related to myelination and oligodendrocyte differentiation among the genes whose expression was upregulated by VL are shown in box-and-whisker diagram. **e**, Expression patterns of oligodendrocyte differentiation marker molecules: OPC, oligodendrocyte precursor cell; pre-OL, pre-oligodendrocyte; OL, mature oligodendrocyte. Genes marked with an asterisk are significantly up-regulated by VL as shown in e. **f**, The box-and-whisker diagram shows the genes classified in GO term “synapse” that are down-regulated by VL exposure.

### Effect of VL on depressive-like behavior in social defeat mice

Activity of the MHb and LHb correlates with serotonin secretion and plays an important role in depression. Increased activity of the LHb releases presynaptic neurotransmitters and causes depressive symptoms^20^. Since OPN5-RGCs are one component of a polysynaptic pathway that projects to the habenula region, we assessed the effect of VL in depression using the social defeat stress (SDS) model. Mice were subjected to SDS for 10 days. Mice susceptible to SDS showed a decreased social interaction (SI) ratio and were selected for further study. SDS-susceptible WT and OPN5 knockout (OPN5 KO) mice were exposed to VL for 3 h per day (ZT9-ZT12) over 10 days, and depression-like behavior was confirmed by SI and other behavioral tests, including sucrose preference (Fig. 5a, b). The data demonstrated that SDS-susceptible WT mice without VL exposure showed significantly reduced SI ratio even after 13 days following the last defeat session. By contrast, VL-exposed WT mice showed increased SI ratio compared to mice under white light, indicating the efficacy of VL as an antidepressant in the improvement of social behavior compared to white light, which does not include VL spectrum (Fig. 5c). VL-exposed OPN5 KO mice, however, exhibit depressive behavior, as indicated by the reduced SI ratio. The total distance travelled between the groups were not altered by light conditions (Fig. 5d). Moreover, WT mice maintained after SDS induction in the absence of VL stimulation showed decreased time in the social interaction zone and increased time in the corner zone away from the ICR mouse (Extended Data Fig. 5a, b). In the sucrose preference test, VL treatment for 10 days increased the sucrose intake in SDS susceptible mice, while SDS decreased the sucrose intake under the white light condition (Fig. 5e). However, in the tail suspension test, no effect of VL could be seen between the three groups (Extended Data Fig. 5c). Together, these behavioral data suggest a novel approach of VL in the treatment of depressive-like behavior. Recent studies have shown the involvement of PVT in the stress paradigm^21,22^. Similarly, many studies reported that the NAc is required for driving social behavior. To investigate the neural mechanism underlying the impact of VL on SDS mice, we examined the role of VL on cFos expression in the brains of defeated mice. In our study, cFos expression in PVT and NAc brain regions in SDS mice subjected to VL was significantly elevated, whereas very few cFos positive cells were found in SDS mice without VL (Fig. 5f, g). The reduced cFos positive cells in defeated mice compared to control was different from previously reported studies that show increased cFos activation in stressed mice^23,24^ (Fig. 5g). Moreover, the activation of cFos were also seen in other parts of the brain including medial amygdala, hypothalamus, piriform cortex, and Lateral septal ventricle (Extended Data Fig. 5d). Together, the results suggest that social defeat stress mice are associated with reduced neural activation, which could modulate the onset of depression.

**Fig. 5.**
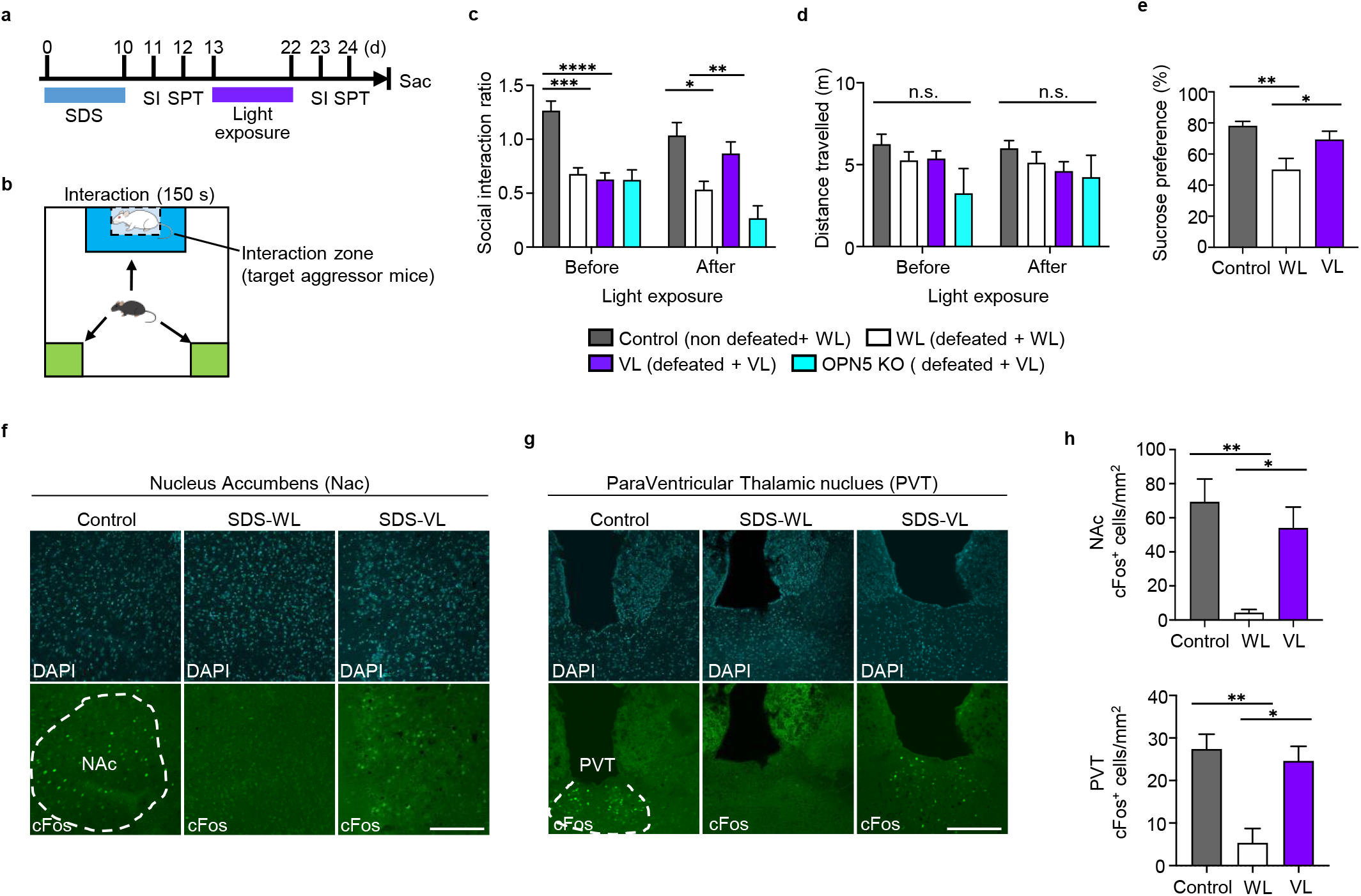
VL treatment promotes social behavior. **a**, Schematic diagram of the SDS model and the schedule of the VL exposure experiment. During the period shown in the figure, mice were exposed to VL for 3 h per day from 5 pm to 8 pm. **b**, Schematic of the SI chamber showing both the interaction zone and corner zone in the social interaction (SI) test. Bar graphs show the social interaction ratio (**c**) and total distance traveled (**d**) in the SI test before and after light exposure. Control (non-defeated + WL) mice (n=18), WL (defeated + WL) mice (n=17), VL (defeated + VL) (n=22), OPN5 KO (defeated + VL) (n=4). *p=0.0421, **p=0.0022, ***p=0.0001, ****p<0.0001, Mixed effect analysis followed by Tukey’ s test. **e**, Sucrose preference test using SDS mice after light exposure. *p = 0.0375, **p < 0.0026, One-way ANOVA analysis followed by Tukey’s test; Control (n=12), WL (n=10) and VL (n=14). **f-g**, Cryosections of mouse brains exposed to white light or VL were immunostained with cFos and DAPI. Left: Nucleus Accumbens, Right: PVT region, Scale bar 50 μm. **h**, Quantitative analysis of cFos positive cells in the depression-related regions. NAc: *p=0.0168, **p=0.0019, PVT: *p=0.0129, **p=0.0057, One-way ANOVA analysis followed by Tukey’s test; Control (n=3), WL (n=3) and VL (n=3). Bar graphs show mean±s.e.m.

## Discussion

We have previously uncovered that VL prevents the progression of myopia which is induced by axial length elongation in chick and human^25^. We found that the thinning of the choroid and the elongation of the eyeball along the visual axis in lens-induced myopia are suppressed by VL^26^. The effect on myopia requires OPN5 in RGCs, indicating that the RGC-brain network through OPN5 is crucial for eye development. OPN5 is the most conserved non-visual opsin in mammals compared to OPN3 and OPN4, and is found in frogs, birds, and mammals^27,28^. Although *Laurasiatheria, Afrotheria, Xenartheria, Euarchontoglire*, and other mammals have different opsins related to visual information depending on their living environment, the fact that all these mammals have OPN4 and OPN5 suggests that OPN4 and OPN5 play a crucial role in controlling tissue functions in response to light environment.

### The role of VL on cognitive function

Here, we have observed that light stimulation by VL improves cognitive function in aged mice and depression-like behavior induced by SDS. It is known that serotonin levels and sleep change in response to the season and daylight hours, affecting the cognitive functions of the brain^29^. In addition, bright light treatment in the early morning improves nonseasonal major depressive disorder, indicating the effects of light on brain function in addition to the phase-resetting effects of circadian rhythms^30^. However, fluorescent lights and white LEDs do not contain VL, and we have not observed the improvement in cognitive function or depression under normal lighting conditions with white LED. The subset of RGCs expressing OPN5 in *Opn5cre; Ai14* mice are largely different from OPN4-positive RGCs indicating that the signal from VL is discriminated from blue light. ipRGCs are classified into six subtypes of RGCs (M1-M6) for rodents and four subtypes (M1-M4) for humans. In response to blue light, *Brn3b* (-) M1 ipRGCs innervates SCN in the hypothalamus to regulate sleep, body temperature and cognitive function^5,31^. Although the changes in hippocampus-dependent cognitive function were associated with the SCN in OPN4, the tracer WGA expressed in OPN5-RGCs indicated that the signal of VL projects to the MHb but not the SCN from RGCs. RNA-seq showed that genes such as *Sox10, Cnp, Mag, Mbp* and *Mobp* were increased in hippocampus after VL stimulation. During aging, the decreased capacity of neuronal progenitor cells (NPCs) and *de novo* myelination impairs memory consolidation, which is defined as a temporal coordination of neural activity among the different brain regions to stabilize working memory^32^. Taken together, VL may have a crucial role for cognitive function by maturation of myelin-forming oligodendrocytes from NPCs. The mechanism to induce oligodendrocyte associated genes via VL remains unknown, and further experiments are required to decipher the underlying mechanism.

### The regulation of mood by VL depends on OPN5

The T7 cycle, in which light and darkness are repeated every 3.5 h, induces depression-like behavior in the sucrose preference test, tail suspension test, and forced-swim test. The depletion of *Brn3b*(+) ipRGCs abolished the T7 cycle-induced depression, and PHb is the central brain region to regulate mood owing to blue light^2^. We revealed that VL improves depression-like behavior by increasing social interaction and the increased sucrose preference, induced by SDS without bright light. Although it remains unknown whether the VL stimulation requires OPN5-RGCs, whole-body OPN5 KO mice indicate that the improvement of depression-like behavior by VL depends on OPN5. Moreover, the tracer of WGA from OPN5-RGCs was observed in MHb but not PHb, indicating that not only the neural circuit but also the mechanism to regulate memory and mood are discriminated. The NAc is the key brain region regulating emotion, addiction, and depression through medium spiny neurons (MSNs) which are GABAergic projection neurons^33^. VL in a depression model increases cFos in NAc as well as the PVT located at the epithalamus. Further experiments will be required to understand the neural circuit from MHb to NAc and PVT after VL stimulation.

### Perspective

Our finding revealed a novel function of VL in regulation of cognitive function and depression. The well-conserved OPN5 may be a non-visual photoreceptor for sensing the light environment and the brain function is maintained by VL during the lifespan. OPN5 is expressed not only in RGCs but also in skin and in the hypothalamic POA in which OPN5 regulates thermogenesis in BAT. Interestingly, retinal progenitor cell-specific depletion of OPN5 using Rx^cre^ had no impact on the regulation of thermogenesis via VL^13^. Although we speculate OPN5-RGCs project to MHb leading to the regulation of cognitive function and mood, the central mechanism sensing VL needs to be addressed. Non-invasive intervention using VL could provide a novel solution for the super-aging society.

## Methods

### Mice

All experiments were performed in accordance with animal rights guidelines of the Japan Ministry of Health, Labor and Welfare for care and use of laboratory animals approved by Animal care and committees of Keio University, School of Medicine. The mice were housed at 22 °C in a 12 h light/12 h dark cycle with free access to food and water ad libitum unless mice were subjected to any treatment. Eight week-old male C57B6/J mice and male ICR mice were purchased from SLC (Shizuoka, Japan) and were used for SDS and for all behavior experiments. *Opn5^cre^; Ai14* mouse line was generously provided by Dr. Richard A. Lang (Cincinnati Children’s Hospital Medical Center, Cincinnati, OH, USA)^14^.

### Light exposure to mice

Mouse cages were placed in a light exposure device (newopto; Kanagawa, Japan) attached with LEDs (White: NF2W757GT-F1, Violet: NSSU123, NICHIA CORPORATION; Tokushima, Japan) on the top of the box. Mice in the control group were housed under white light (475 lux) from 8 am to 8 pm in a 12 h light/12 h dark cycle. Mice in the experimental group were also exposed to white light under the same conditions except for the violet light exposure (670 μW/cm^2^). In the learning and memory experiments, aged mice were exposed to violet light from 10 am to 12 pm daily for the indicated period. In the depression experiments, SDS mice were exposed to violet light from 5 pm to 8 pm for 10 days.

### Contextual fear conditioning test

On day 1, mice were placed into an experimental box (W 19 cm x D 19 cm x H 28 cm) and allowed to explore freely for 180 s followed by 1.0 mA electric shock for 2 s. Another 1.0 mA shock for 2 s was given after 30 s and immediate freezing was measured every 10 s by a visual count for 60 s. Following the experiment, the mice were returned to their home cage. Contextual freezing without a tone was assessed for 180 s, 24 h after the shock, counting freezing every 10 s. During each session, the maze was wiped with 70% EtOH and with 1% acetate to remove the odor of the mouse in previous trial. Freezing was counted by two experimenters and averaged for analysis.

### Barnes maze test

The maze consists of a circular and white platform (100 cm in diameter) with 20 equally spaced holes (5 cm in diameter) around the edge of the platform, elevated 90 cm above the floor. For visual cues, four pictures with different colors and shapes are placed around the maze. In the adaptation period, a mouse was placed in a white cubical start chamber in the center of the maze, then gently guided to a small chamber termed a target hole. The adaptation trial was repeated 3 times. During the spatial acquisition period, the mouse was allowed to explore the maze for 3 min. If the mouse entered the target hole, spent time around the target hole for 30 s, or it passed for 3 min, the mouse was allowed to stay for 1 min in the hole. After each trial, the mouse was placed in its home cage until the next trial. The trial was repeated 3 times/day for 6 days. Probe trials were conducted to assess the short-term and long-term memory retention with the target hole covered with a lid. The mouse was allowed to explore the maze for 90 s. During each session, the maze was wiped with 70% EtOH and with 1% acetate to remove the odor of the mouse in the previous trial. The data were acquired by video-tracking software, Anymaze (Stoelting; Illinois, USA).

### AAV production and injection in mouse retinal cells

pAAV-DIO (double-floxed inverted open reading frame)-WGA vector was purchased from VectorBuilder (Chicago, USA). AAV-DIO-WGA was generated by co-transfection of pAAV-DIO-WGA, pAAV-RC and pAAV-Helper plasmids into 293T cells using PEI MAX (Polysciences; Pennsylvania, USA). 72 h after transfection, cells and medium were collected and AAV particles were purified by AAVpro Purification Kit (Takara Bio; Shiga, Japan) according to the manufacturer’s instructions.

For retinal injection of AAV particles, mice were anaesthetized with a combination of midazolam (Sandoz; Holzkirchen, Germany), medetomidine (Orion; Espoo, Finland), and butorphanol tartrate (Meiji Seika Pharma; Tokyo, Japan). AAV-DIO-WGA particles were bilaterally injected into the retina using a microsyringe and needle (ITO; Shizuoka, Japan). After 6-8 weeks, mice were anaesthetized and perfused with 25 ml PBS followed by 20 ml of 4% paraformaldehyde (PFA) solution. Brains and eyes were dissected and postfixed in 4% PFA for 24 h for immunohistochemistry.

### Social defeat stress (SDS)

SDS was performed as previously described^34^. Briefly, B6 mice were introduced to the home cage of ICR and allowed the defeat for 10 min. After 10 min physical defeat, B6 mice were separated by a perplexing glass divider from their respective ICR, allowing sensory contact for 24 h. The defeat process was carried out for 10 consecutive days with novel ICR mice each day. After the last defeat session, all B6 mice were then returned to a single cage and proceeded for behavioral experiments.

#### Social interaction test

Social interaction is used to assess the avoidance behavior as one of the criteria for depressive-like phenotype. The defeated and control mice were tested the next day after the last day of a defeat session. The test chamber consists of two zones, the interaction zone (15 cm × 30 cm) rectangular area projecting 8 cm around the wire-mesh enclosure and the corner zones (10 cm × 10 cm). Briefly, in the first step (“habituation”), defeated mice were placed in the open chamber and set free to explore the chamber for 150 s without target (aggressor ICR mice) in the open field box. Next, 30 s intervals where the mice were returned to their home cage and an unfamiliar target mouse was placed into the cage inside the cylinder. Finally, in the last step (“interaction”), defeated mice were reintroduced to the chamber where the target is kept inside the wired cage for another 150 s. The data were acquired by video tracking software ANYmaze. The social interaction ratio and time spent by each mouse in each zone were quantified by using GraphdPrism8.

##### Sucrose Preference Test (SPT)

Mice were subjected to two identical bottles, one with 1% sucrose solution and other with regular tap water for 48 h as a habituation period. To avoid any bottle preference, the bottles’ positions were switched every 24 h during the habituation period. After 48 h of habituation period, mice were deprived from food and water for 12 h before the actual testing. During the testing period mice were given two identical bottles, each filled with 1% sucrose solution and regular tap water for 12 h. Finally, the weight of the bottles before and after the test was recorded, the percentage of sucrose preference in reference to the total liquid consumption was calculated using the equation: Sucrose preference (%) = sucrose intake/ (sucrose intake + water intake) x 100

##### Tail Suspension Test (TST)

The tail suspension test was performed on 12^th^ day of the social defeat stress experiment. Briefly, mice were suspended by placing one end of tape on the tail and the free end of the tape on a suspension bar in a way that it does not obstruct the camera view since the entire TST session will be scored, and obstructing the camera will lead to the inability to assess behaviors during that time. Each session lasted for 6 min, and at the end of the session, each mouse was returned to their home cage and the tape gently removed from the tail. After returning the mice to their colony room, the fecal boli and urine from the collection trays were discarded and the apparatus wiped with a sterilizing solution before the start of another session. Immobility time (%) was calculated by the time mice were immobile divided by total time.

##### Immunohistochemistry

Animals were transcardially perfused with 25 ml PBS followed by 20 ml of 4% paraformaldehyde solution. Brains and eyes were then removed and soaked in 4% paraformaldehyde for 24 h, followed by gradually to 10% sucrose solution overnight, next 20% sucrose and finally 30% sucrose solution for 2-3 days, and embedded in OCT-compound (Sakura Finetek; Tokyo, Japan). Each brain and retina were sliced into a series of consecutive 20-30 μm coronal sections on a cryostat and placed on an APA glass slide. Brain and retina sections were first incubated for 30 min with blocking solution (1% BSA, 0.25% Triton X in PBS). Sections were then incubated for 1 day at 4 °C in blocking solution with following antibodies: rabbit anti-cFos (226003, Synaptic Systems; Goettingen, GERMANY, 1:1000); goat anti-Wheat Germ Agglutinin (WGA) (AS-2024, VECTOR; California, USA, 1:1000); and mouse anti-GAD65 (ab26113, abcam; Cambridge, UK, 1:1000). Sections were rinsed in PBS 3 times for 5 min and then incubated for 2 h at RT in blocking solution with the corresponding Alexa-conjugated secondary antibodies (Thermo Fisher Scientific; Massachusetts, USA). Finally, sections were rinsed in PBS for 5 min, 3 times and mounted by cover slips with ProLong™Glass Antifade Mountant with NucBlue™ Stain (Thermo Fisher Scientific). For retinal whole mount images, retinas were extracted from the eye after fixation and several slits were made to mount. The retinas were incubated for 1 day with blocking solution (1% BSA, 0.5% Triton X in PBS). The retinas were then incubated for 2-3 days at 4 °C in blocking solution with following antibodies: rabbit anti-RBPMS (1830-RBPMS, Phospho Solutions; Colorado, USA, 1:1000); rabbit anti-Brn3a (411003, Synaptic Systems, 1:1000); rabbit anti-Melanopsin (AB-N39, Advanced targeting systems; California, USA, 1:1000); and goat anti-WGA (1:1000). The retinas were rinsed in PBS 4 times for 30 min and then incubated for 1 day at 4 °C in blocking solution with the corresponding Alexa Fluor-conjugated secondary antibodies. Finally, sections were rinsed in PBS 4 times for 30 min and mounted using ProLong™Glass Antifade Mountant (Thermo Fisher Scientific). All images were acquired by FV3000 confocal microscopy (Olympus; Tokyo, Japan). For image quantification 20x magnification used for all the sections for quantitative analysis. The cFos positive cells were counted using three sections per mice in ImageJ.

##### Tissue Preparation

After the last behavioral tests, mice were euthanized by cervical dislocation. The entire brain of each mouse from aged, young, and SDS were removed. The tissues were collected in two halves, one half were used for immunostaining and the other half for gene expression analysis. All the tissues (hippocampus, cortex) were quickly frozen in liquid nitrogen and then stored at −80 °C until further analysis.

##### Total RNA Extraction

Total RNA was extracted from hippocampus tissue. The tissue was homogenized in QIAzol lysis reagent (Qiagen, Germany), and total RNA was extracted using RNeasy mini kit (Qiagen, Germany). All RNA samples quality and quantity were determined using RNA Nano kit (2100 Agilent Bioanalyzer Technologies, CA, USA). Only samples with an RNA integrity number greater than 8.0 were used for gene expression analysis.

#### Quantitative real time PCR (qRT-PCR)

Total RNA was isolated from each tissue in QIAzol lysis reagent (Qiagen, Germany) and the concentration was measured by Nanodrop One^c^ (Thermo Scientific). 500 ng of total RNA was used for cDNA synthesis by reverse transcriptase RNA kit (Applied biosystems, Japan). Quantitative real-time PCR was performed according to the protocol described in our previous report. Briefly, the PCR reaction using SYBR Green, was conducted with a Quant studio 5 under the following conditions: 2 min at 50 °C followed by 40 cycles at 95 °C for 10 s and 60 °C for 1 min. Each transcript value was calculated as the average of duplicate samples across experimental conditions. Values were normalized to GAPDH as an internal control. The data were analyzed by comparing the cycle threshold (Ct) values of the different groups with the 2^−ΔΔCt^ method.

##### Library Preparation

Four mice from each group (WL and VL) were chosen randomly for RNA-seq analysis. Hippocampus RNA-seq libraries were prepared in accordance with New England Biolab protocols (NEBNext_ Ultra II RNA library Prep kit for Illumina, NEB, USA). Briefly, rRNA were depleted from 1 μg of total RNA using the NEBNext rRNA depletion kit (Huma/Mouse) and ribo-depleted RNA was quantified by Quant-iT ™ RiboGreen ™ RNA Assay Kit (Invitrogen). Using a reverse transcriptase and random primers, first- and second strand cDNAs were synthesized. The cDNA was subjected to an end repair reaction followed by adapter ligations to the end of the DNA fragment. Size selection of DNA fragments was performed by means of SPRISelect Beads (Beckman Coulter, Brea CA, USA) and then PCR enrichment of the adapter-ligated library was conducted (12 cycles of PCR). The size and quantity of the library were verified using HS DNA assay (2100 Agilent Bioanalyzer, USA), and KAPA qPCR. The libraries were sequenced by HiSeqX illumina

## Acknowledgements

This study was supported by Tsubota Laboratory, Inc. and Sumitomo Dainippon Pharma Co., Ltd. This study was also funded by Program for the Advancement of Research in Core Projects, Keio University. We thank Core Facility, Collaborative Research Resources, Keio University School of Medicine for technical support. Also, we would like to thank Sarah Y. Robertson for reviewing and providing valuable feedback on this article.

## Author contributions

N. S., P. G., M. H. and K. T. conceptualized and designed the experiments. N. S., N. T., and K. O. performed behavioral experiments of aged mice. N. S. and R. T. carried out tracing experiments. T. S. contributed to bioinformatic analysis of RNA-seq data. Y. H., R. T., K. O., and H. O. carried out the immunohistochemical studies and gene expression analysis. N. S., P. G., and K. O. conducted SDS mice experiments. N. S., P. G. and M. H. wrote the manuscript. N. B., Y. M., R. A. L, M. M. and K. T. critically reviewed the manuscript.

