## Extended data Fig.1-5 for "Violet light modulates the central nervous system to regulate memory and mood"

a

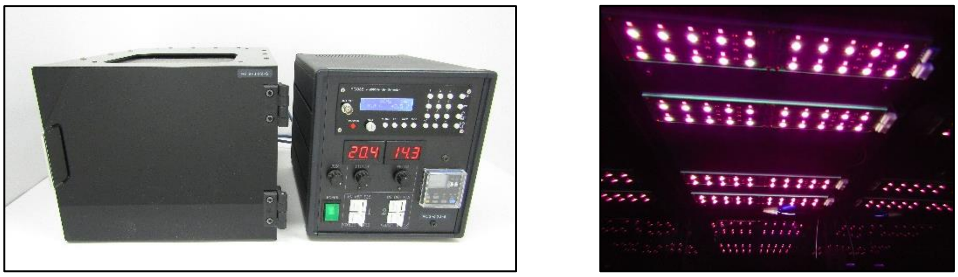

b

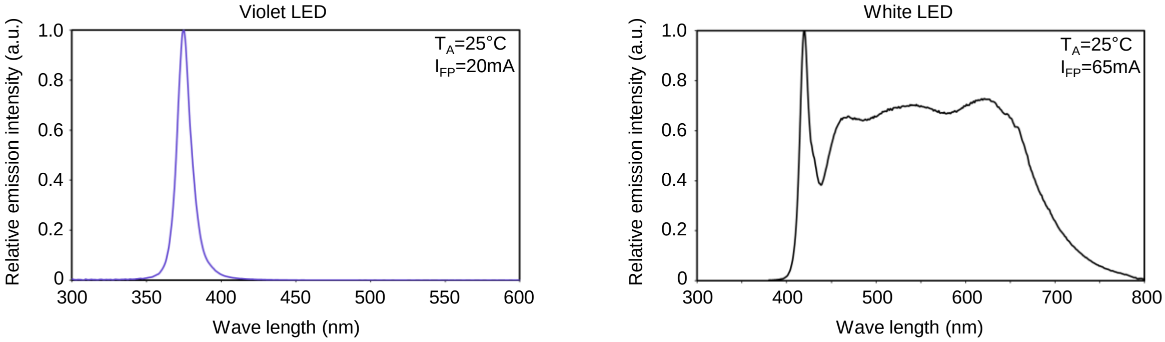


**Extended Data Fig. 1| Light-exposure device and *Opn5* gene expression in young and aged mice retina**

**a**, (left) Light-exposure device (right) A view of violet light irradiation. **b**, The spectrum of the LED attached to the device, violet (left) and white (right).

a

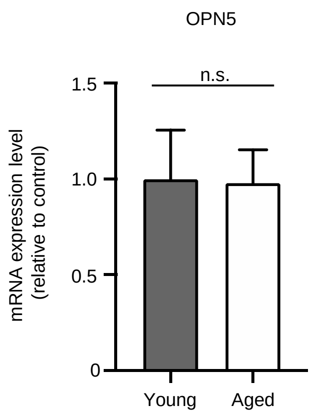


**Extended Data Fig. 2| *Opn5* gene expression in young and aged mice retina**
**a**, Expression of *Opn5* gene in young (8 weeks old, n=5) and aged (89 weeks old, n=4) mice retina. Expression levels of each gene were quantified by RT-PCR. *Gapdh* was used as an internal control. Bar graphs show mean±s.e.m.

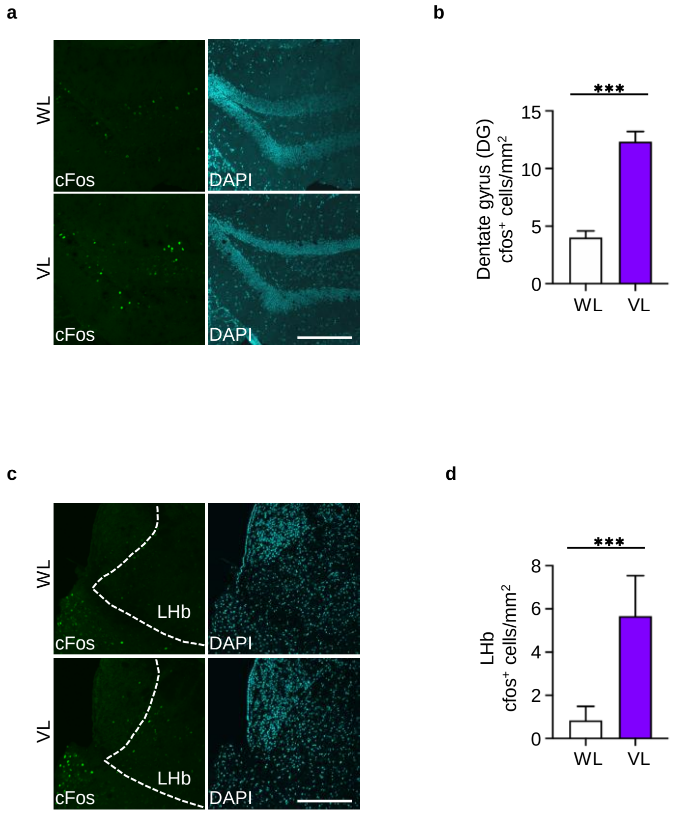


**Extended Data Fig. 3| cFos expression in VL-exposed mouse brain**

**a**, Expression of cFos in the dentate gyrus. Frozen sections of mouse brains irradiated with white light (upper panel) or VL (lower panel) were immunostained with cFos and DAPI. **b,** Quantitative analysis of cFos positive cells, Student's t-test, ***p=0.0001, **c**, Expression of cFos in the LHb. Frozen sections of mouse brains irradiated with white light (upper panel) or VL (lower panel) were immunostained with cFos and DAPI. **d,** Quantitative analysis of cFos positive cells. ***p=0.0001, Student's t-test; WL (n=3) and VL (n=3). Scale bar 50 µm. Bar graphs show mean±s.e.m.

**a**

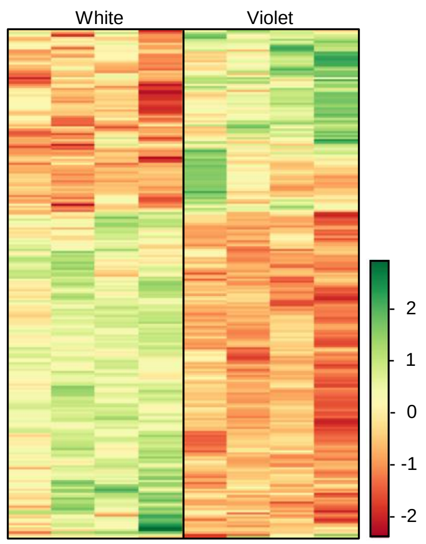


**Extended Data Fig. 4| VL modulates gene expression in hippocampus**

**a**, Differentially expressed genes in the hippocampus of VL-exposed mice are shown in the heatmap (FDR<0.1). Up- and down-regulated genes are shown in green and red, respectively.

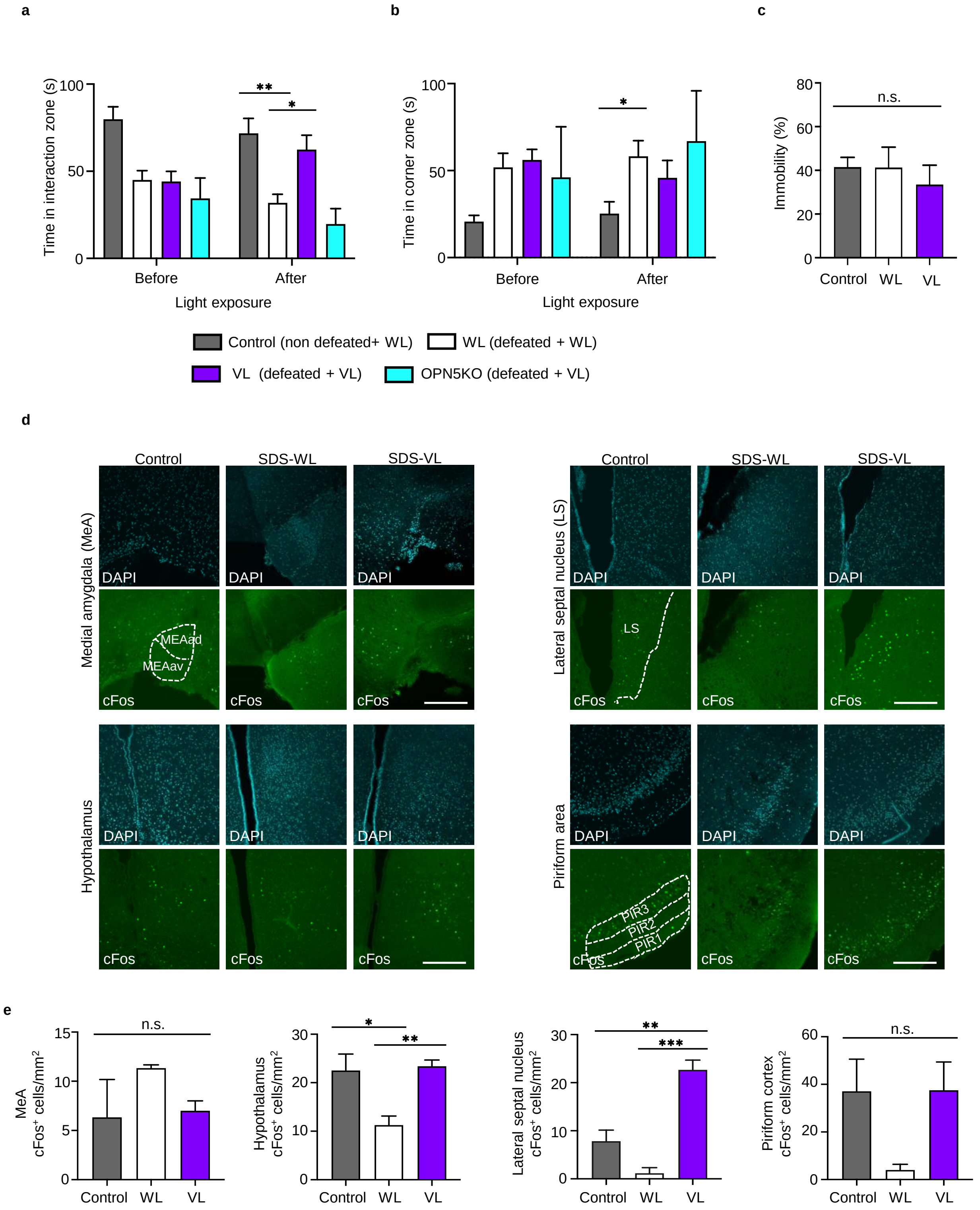


**Extended data Fig. 5| VL treatment promotes social behavior**

**a**, Total time spent in the interaction zone when there is a ICR mouse present. **b**, Total time spent in the corner zone when there is a ICR mouse present. **c,** Immobility time in the TST. Control (non-defeated + WL) mice (n=18), WL (defeated + WL) mice (n=17), VL (defeated + VL) (n=22), OPN5 KO (defeated + VL) (n=4). *p=0.0417, **p=0.0039, Mixed effects analysis followed by Tukey' s test. **d**, Cryosections of mouse brains exposed to white light or VL were immunostained with cFos and DAPI. Upper Left: Medial amygdala, Upper Right: Lateral septal nucleus, Lower Left: Hypothalamus, Lower Left: Piriform cortex, Scale bar 50 μm. **e**, Quantitative analysis of cFos positive cells. MeA: ns, LS: **p=0.0036, ***p=0.0005, Hypothalamus: *p=0.0161, **p=0.0075, Piriform cortex: ns, One-way ANOVA analysis followed by Tukey's test; Control (n=3), WL (n=3) and VL (n=3). Bar graphs show mean±s.e.m.
